# Biological control of chili damping off disease, caused by *Pythium myriotylum*

**DOI:** 10.1101/2020.07.28.224519

**Authors:** Sajjad Hyder, Amjad Shahzad Gondal, Zarrin Fatima Rizvi, Muhammad Irtaza Sajjad Haider, Muhammad Inam-ul-Haq

## Abstract

*Pythium myriotylum* is a notorious soil-borne oomycete causes post-emergence damping off in chilli pepper. Of various disease management strategies, utilization of plant growth promoting rhizobacteria (PGPR) in disease suppression and plant growth promotion is eye catching strategy. The present study was performed to isolate and characterize PGPR indigenous to chili rhizosphere in Pakistan, and to test their potential to suppress damping off and plant growth promotion in chilli. Out of total 28 antagonists, 8 bacterial isolates (4a2, JHL-8, JHL-12, 1C2, RH-24, 1D, 5C and RH-87) significantly suppressed the colony growth of *P. myriotylum* in dual culture experiment. All the tested bacterial isolates were characterized for biochemical attributes, and 16S rRNA sequence based phylogenetic analysis identified these isolates as *Flavobacterium* spp., *Bacillus megaterium, Pseudomonas putida, Bacillus cereus* and *Pseudomonas libanensis*. All the tested bacterial isolates showed positive test results for ammonia production, starch hydrolase (except 4a2), and hydrogen cyanide production (except 4a2 and 1D). All the tested antagonists produced indole-3-acetic acid (13.4-39.0 μg ml−1), solubilized inorganic phosphate (75–103 μgml^-1^) and produced siderophores (17.1–23.7%) in vitro. All the tested bacterial isolates showed varied level of susceptibility and resistance response against different antibiotics and all these bacterial isolates were found non-pathogenic to chill seeds and notably enhanced percentage seed germination, plumule, redical length and vigor index over un-inoculated control. Additionally, under pathogen pressure, bacterization increased the defense related enzymes (PO, PPO and PAL) activates. Moreover, chilli seeds treatment with these bacterial isolates significantly suppressed damping-off caused by *P. myriotylum*, and improved PGP traits as compared to control. In addition, a positive correlation was noticed between shoot, root length and dry shoot and root weigh and a negative correlation was seen between dry shoot, root weight and seedling percentage mortality. These results showed that native PGPR possess multiple traits beneficial to the chilli plants and can be used to develop eco-friendly and effective seed treatment formulation as an alternative to synthetic chemical fungicides.

## INTRODUCTION

Chilli pepper (*Capsicum annuum* L.) a member of *Solanaceae* family is an important vegetable crop, stands forth worldwide and particularly cultivated in Asia on large scale (Tariq et al., 2014). It covers almost 20% of the total vegetable grown area in Pakistan. Chilli pepper is consumed as fresh or processed spice, and serves as a good source of vitamin A and C, phenolics and carotenoids. Capsaicinoid compound derived from chilli has many ethnopharmacological applications including anticancer, anti-obesity treatment, temperature regulation, pain therapy, and as antioxidant (Meghvansi et al., 2010). Chili crop is vulnerable to more than 100 different types of pathogens during various growth stages (Jayapala et al., 2019). Of different microbial diseases, damping off and root rot disease caused by *Pythium myriotylum* Drechsler., is the most devastating disease in terms of seedling mortality at very early growth stages in nurseries when the seedlings are in cotyledonous stage. *Pythium* spp. are disease causative fungal-like organisms that result in 90% plant death as pre and/or post-emergence damping off under favorable conditions. A study has shown that damping-off may effect from 5 to 80% of the seedlings and result into huge economic losses to the farmers (Lamichhane et al., 2017). This disease is characterized by typical symptoms of rotten roots, necrosis, wilt, water soaking lesions, and decay of young seedlings (Horst, 2013). *P. myriotylum* is one of the most commonly occurring species in greenhouse, warm and moist soil and present wide host range (Ben-Yephet and Nelson, 1999). Watery soaked, sunken lesion can be seen on the stem at soil level or below the soil on roots, causing the seedling to fall over the ground (Smith Jr, 1975) and excessive soil moisture favors the development and movement of zoospores which attack the host plants.

Of various other practices, chemical seed coating is widely adopted in agriculture to control the disease (Dorrance et al., 2009; Kandel et al., 2016; Rothrock et al., 2012). Chemicals such as bleach, hydrogen peroxide, ethanol, and fungicides are extensively used to kill pathogen inoculum present on seed coats (Mancini and Romanazzi, 2014). Chemical seed treatment is an effective practice in controlling the soil and seed borne pathogens but can pose detrimental effect on seed germination and cause phytotoxicity (du Toit, 2004). Besides this, pesticide residues in soil and water are potential threat to human and environment (Lamichhane et al., 2017; Ouyang et al., 2016), and many of these chemicals are declared as carcinogen pollutants in many countries (Bressa et al., 1997). Non-judicial use of many of the synthetic pesticides and fungicides has come under increasing public scrutiny in different countries (Bourguet and Guillemaud, 2016) and reports on pest resistance development are also increasing the threat for farmers (Onstad, 2013). Furthermore, these fungicides are noxious to the survival of beneficial rhizosphere microbes (Hussain et al., 2009). Thus, there is strong need to find cost effective and environmentally safe alternatives which can minimize or eliminate the dependency on synthetic pesticides.

PGPRs are free-living or plant root colonized bacteria that confer plant growth promotion (Glick, 2012) and do not cause any harm to their hosts (Ryan et al., 2008). Bacterial belonging to *Pseudomonas, Azospirillum, Azotobacter, Klebsiella, Enterobacter, Alcaligenes, Arthobacter, Burkholderia, Bacillus*, and *Serratia* spp. improve plant growth (Kloepper et al., 1989; Souza et al., 2015), used as biocontrol agents (Chen et al., 2009; El-Sayed et al., 2014; Labuschagne et al., 2010; Liu et al., 2007) and biofertilizers (Vessey, 2003). PGPR promote plant growth by different mechanisms which include the production of Indole acetic acid (IAA) (Etesami, et al., 2015), phosphate solubilization (Panhwar et al., 2014), atmospheric nitrogen fixation (Kuan et al., 2016), ACC deaminase activity (Chen et al., 2013) and zinc solubilization (Gupta et al., 2015). PGPRs suppress plant pathogens by employing various mechanisms such as competition, siderophores production, antagonism and induced systemic resistance (Gómez-Lama Cabanás et al., 2014) which activates multiple defense-related enzymes to challenge the against a broad spectrum of phytopathogens (Cazorla et al., 2007; Vanitha and Umesha, 2011). Peroxidases (PO) have been involved in many defense-related mechanisms, including the hypersensitive reaction, lignification, cross-linking of phenolics and glycoproteins and the production of suberization and phytoalexin (Wojtaszek, 1997). Polyphenol oxidase (PPO) catalyzes the oxidation of phenolics to free radicals that reacts with biological molecules, thus hindering the pathogen development (Jockusch, 1966). Phenylalanine ammonia-lyase (PAL) plays an important role in regulation of phenylpropanoid production (Achnine et al., 2004) and synthesis of various defense-related secondary compounds such as phenols and lignin (Tahsili et al., 2014).

Biological control is an alternative strategy to reduce the dependency on agro-chemicals in crop disease management programs (Postma et al., 2003) and use of PGPR in disease management is helpful to reduce the detrimental effects of agro-chemicals on the environment. Many reports are available on the biocontrol potential of PGPR against *Pythium* spp. and plant growth promotion effect in tomato (Al-Hussini et al., 2019), potato (Kenawy et al., 2019), cucumber (El-Tarabily et al., 2009), sugar beet (Williams and Asher, 1996), cereals (Labuschagne et al., 2010) and many other major crops. In many cases, it was observed that imported bioformulations sometimes fail to act up to their maximum potential due to climate change (Compant et al., 2010), nutrient availability (Kandeler et al., 2006) and rhizosphere competence of the microbes (Lugtenberg and Kamilova, 2009). Thus, the identification and characterization of PGPRs indigenous to chilli rhizospheres is important to screen bacterial isolates that can suppress *P. myriotylum* inoculum and enhance chilli growth in nurseries and greenhouses.

Keeping in consideration the importance of chilli production in eco-friendly environment, the aim of this study was to isolate and screen native rhizobacteria for their biocontrol potential against most virulent strain of *P. myriotylum* in vitro, to characterize the bacterial agents on the basis of morphological characters and also by 16S rRNA sequence analysis, to study the effect of bacterial treatment on seed germination, and to study PGPR ability to suppress *P. myriotylum* – induced damping-off and PGP effects on chilli in pot experiments under growth room conditions.

To our knowledge, this is the first report of using native PGPRs suppressing *P. myriotylum* and enhancing growth promotion in chilli from Pakistan.

## MATERIALS AND METHODS

### Pathogen inoculum

The strains of *Pythium myriotylum* D., (PMyr-1 and PMyr-2) were previously reported as the causal agent of damping-off and root rot in chili pepper (*Capsicum annum* L.) from Punjab, Pakistan (Hyder et al., 2018). Pathogen was identified on morphological and molecular basis. The ITS1 and ITS2 rDNA sequences of these two virulent strains had been submitted in GenBank database (accessions no. MF143429 and MF143430), which displayed 99% identity with of *P. myriotylum* (accession no. HQ643704).

### Sampling and isolation of bacterial isolates

Major chilli growing fields in Rawalpindi (33.5651° N, 73.0169° E) Punjab Pakistan were surveyed, and rhizospheric soil samples strictly adhering to chilli plant roots were taken from 15 to 20 cm depth along with the plant roots. All the soil samples were immediately processed for the isolation of rhizobacteria after reaching the laboratory. Bacteria were isolated from 10g of soil samples by serial dilution plating on nutrient agar (NA) (HiMedia Laboratories) medium containing Petri plates (Joseph et al., 2012). For the isolation of root colonizing bacteria, 1g of the root samples were washed with tap water, surface sterilized using 70% ethanol for 5 min followed by 1% sodium hypochlorite (NaHOCl) for 2 minutes and then washed five times with sterilized distilled water (Kuan et al., 2016). Sterilized roots were crushed in distilled water aseptically with sterilized mortar and pestle, and were streaked on NA medium followed by incubation at 26±2 °C for 24-48 hours. Morphologically discrete bacterial colonies were picked aseptically using sterilized loop and sub-cultured on freshly prepared NA medium containing Petri plates. Bacterial isolates were stored at -80°C in equal volumes of nutrient broth (NB) medium and 30% glycerol for further use in experiments.

### In vitro screening of bacterial isolates against *Pythium myriotylum*

The rhizobacterial isolates (n=110) were tested in repeated experiments for antagonistic potential against two virulent strains of *P. myriotylum* (PMyr-1 and PMyr-2) by using dual culture technique (Rabindran and Vidhyasekaran, 1996) on PDA medium containing Petri plates. Small discs of actively growing *P. myriotylum* (5mm) were placed in the middle of 9cm Petri plates, counter streaked on two sides by each rhizobacterial isolate about 2.5cm from the fungal discs. The control plates contained fungal plugs without the bacterial streaks. Petri plates were incubated at 26±2 °C, and inhibition zones (cm) were measured against each isolate 48 and 96 hours after incubation. Each of the bacterial isolate was tested in five repeats to confirm the results. The percentage mycelial growth inhibition was recorded by using the following formula: 

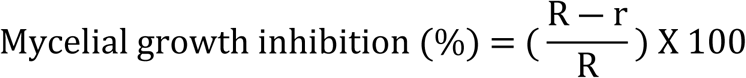

R is radius of fungal mycelial growth in control

r is the radius of fungal mycelial growth in the treatment

### Biochemical featuring of rhizobacterial isolates

Bacterial isolates displaying consistent antagonistic responses in repeated dual culture tests were characterized on the basis of Gram type reaction and fluorescence emission using the standard methods as described earlier (Cappuccino and Sherman, 2005). Potassium hydroxide (KOH) solubility tests were performed using the protocol as previously described by (Kirsop and Doyle, 1991). In this test, 24 hour old bacterial colony grown on NA medium was mixed thoroughly with 3% KOH solution on a glass slide and mixed thoroughly. Formation of mucoid thread confirmed the positive results for the bacterial isolates. Catalase tests were performed in accordance with (Hayward, 1960). In particular, freshly grown bacterial culture on NA medium was mixed with one drop of 3% H_2_O_2_ on a glass slide. Rapid gas bubbles formation confirmed the positive test results. Levan production was tested using the procedure described by (Lelliott and Stead, 1987). Carbohydrate fermentation test was performed in accordance with the procedure previously described by (Aneja, 2001). In this test, overnight grown bacterial cultures were inoculated in screw-capped tubes containing sterilized phenol red carbohydrate fermentation broth (1g Trypticase; 0.5g Sodium Chloride; 0.02mg Phenol red and 0.5g carbohydrate in 100ml of distilled water). Change in medium color from red to yellow indicated the positive test results. Hydrogen sulphide (H_2_S) production test was performed following the protocol described earlier by (Warren et al., 2005). In particular, 24 hours old bacterial cultures grown on NB medium were aseptically inoculated on sulphite indole motility (SIM) medium (HiMedia Laboratories, India) containing tubes followed by incubation at 37 °C. Development of ferrous sulfide (black ppt.) confirmed the positive test results. Oxidase tests were carried out as described by (Hayward, 1960). In this test, 24 hours old bacterial culture was mixed with few drops of 1% of N,N,N′,N′-tetramethyl-p-phenylenediamine (TMPD) solution (Sigma-Aldrich, USA) on Whatman No. 1 filter paper. Appearance of dark purple color within 30 seconds confirmed the positive test results. The test for oxidative fermentative was performed as described by (Hugh and Leifson, 1953), while nitrate reduction and gelatin hydrolysis assays were carried out using the protocol previously used by (Thankamani and Dev, 2011).

### Molecular characterization of rhizobacterial isolates

Bacterial agents displaying promising antagonistic activity were identified using 16S rRNA gene sequencing (Kumar et al., 2015). Total genomic DNA was extracted from bacterial isolates, by using the GeneJet Genomic DNA purification Kit (Thermo scientific Waltham, USA) following the manufacturer’s instructions. The 16S rRNA region was amplified in polymerase chain reaction (PCR) using primer pair 27F [5′ -AGAGTTTGATC-MTGGCTCAG-3′] and 1492R [5′ -GGTTACCTTGTTAC-GACTT-3′] respectively, in 50µl reactions consisting 25–150ng of DNA template, 1X of Taq buffer (10mM Tris pH 9, 50mM KCl, 0.01% gelatin), 200μM of each dNTP, 1.25mM of MgCl_2_, 0.4μM of each primer, and 0.5U of Taq DNA polymerase (Qiagen, Germany). PCR conditions were: initial denaturation of DNA template at 95 °C for 1 minute per cycle, 35 cycles of denaturation at 95 °C for 15 s, annealing at 55 °C for 15 s, extension at 72 °C for 1 minute and a final elongation at 72 °C for 7 minute. Amplified DNA products were then run on 1% (w/v) agarose gel and visualized under UV transilluminator after staining with Ethidium bromide (EB). PCR products (1.5Kb) of 16S rRNA gene were cleaned with Gel and PCR Clean-Up system (Promega, USA), and quantified by NenoDrop. The amplified DNA products were then sent for sequencing to the Department of Crop Sciences, University of Illinois, Urbana, IL, USA. Frequents were sequenced using 27F and 1492R primers, and obtained sequences were joined by Bioinformatics software for life science (DNASTAR software). Sequences were run in the BLAST program (https://blast.ncbi.nlm.nih.gov/Blast) at National Center for Biotechnology Information (NCBI) server to search the closely related sequences. All the retrieved sequences along with tested bacterial isolates sequences were aligned together using CLUSTAL W Program. Evolutionary relatedness between tested bacterial sequences and retrieved sequences was determined by constructing phylogenetic tree using neighbor-Joining (N-J) method in Molecular Evolutionary Genetics Analysis software MEGA X version 10.1.7 with 1000 bootstrap replicates. The evolutionary distances were calculated using Kumara 2-parameter model (K2 + G) (Kumar et al., 2018). The 16S rRNA gene sequences were deposited in the GenBank nucleotide database and accession numbers were obtained.

## BIOASSAYS FOR PLANT GROWTH PROMOTION (PGP) TRAITS

### Ammonia (NH_3_) production

The production of NH_3_ was testes in accordance with (Cappuccino and Sherman, 2005). In particular, twenty-four hour old each bacterial isolates (100μl) grown on nutrient broth medium was inoculated on test tubes containing peptone water (10.0g peptone; 5.0g NaCl; 1000ml distilled water; 7.0 pH) and incubated at 28°C for 48-72 hours. 500µl of Nessler’s reagent (Fisher®, USA) was added to each test tube. Brown to yellow color development confirmed the NH_3_ production.

### Starch hydrolysis

Starch hydrolysis test was performed using the protocol as previously described (Marten et al., 2000). In particular, 24 hours old bacteria were cultured on LB agar medium containing Petri plates amended with 2% starch and incubated at 30±2°C for 48–72 hours. Plates were then flooded with Lugol’s solution. Clear halo zone formation around the bacterial growth confirmed the positive test results of starch hydrolysis.

### Phosphate solubilization

Phosphate solubilizing ability was assessed by following the procedure as previously reported (Verma et al., 2001). For this test, the bacterial cultures were streaked on Pikovskaya’s agar medium (HiMedia laboratories, India) supplied with tricalcium phosphate in Petri plates. Plates were then incubated 28±2°C for 72-96 hours. Formation of clear halo zones encircling the bacterial colonies indicated phosphate solubilization. Phosphate solubilization was quantified by Phosphomolybdate blue color assay as previously described by (Murphy and Riley, 1962).

### Hydrogen cyanide

Production of HCN was assessed by adopting the procedure reported by (Lorck, 1948). In this test, 24 hours old bacterial strains were streaked on NA medium containing Petri plates amended with glycine (4.4gl^-1^). Agar medium was covered with Whatman number 1 filter paper previously dipped in the solution of 0.5% picric acid and 2% sodium carbonate (w/v). Petri plates were Parafilmed, and were incubated at 28±2°C for 96 hours. Development of orange or red color indicated HCN production.

### Indole acetic acid (IAA) detection and quantification

Bacterial isolates were inoculated on LB medium amended with 0.5mgl^-1^ tryptophan/ml, incubated at 28±2°C for 5 days and centrifuged at 3,000 rpm for 30 minutes. The supernatant (2ml of the aliquot) was added with two drops of orthophosphoric acid and 4ml of Salkowaskis reagent (150ml concentrated H_2_SO_4_, 250ml distilled water, 7.5ml 0.5M FeCl_3_.6H_2_O), and incubated at room temperature in dark for 20 minutes (Gordon and Weber, 1951). Development of a pink-red color indicated the production of IAA. Absorbance of IAA was recorded at 530 nm using spectrophotometer (Thermo Scientific, USA) and the concentration of IAA was measured against a standard curve developed from pure IAA solution. There were three replications for each bacterial isolates, and all the mean values were statistically analyzed.

### Siderophores production

Siderophores production was tested in accordance with (Schwyn and Neilands, 1987). In particular, siderophores production test was performed by culturing bacterial isolates (10^8^ cfu ml^-1^) on Chrome azurol S agar medium followed by incubation at 28±2 °C for 72 hours. Change in the color from yellow to orange was an indication of siderophores production. Siderophores production was quantified by CAS-liquid assay (Payne, 1994); Optical density (OD) was measured at 630nm against a reference consisting of CAS reagent. Siderophore contents were calculated by the formula: 

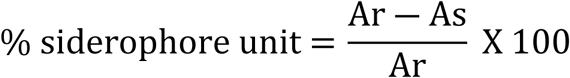

Ar = absorbance of the standard at 630 nm.

As = absorbance of the sample at 630 nm.

### Multiple antibiotic resistance of rhizobacterial isolates

Multiple antibiotic resistance tests were performed to check the level of susceptibility and resistance of rhizobacterial isolates by following the methodology previously described (Singh et al., 2013). The test was performed to screen the bacterial isolates against streptomyces, ampicillin, rifampicin, penicillin G and vancomycin at different concentration levels (0 ppm, 100 ppm, 200 ppm, 300 ppm, 400 ppm and 500 ppm) in vitro. For this, 100μl of 24 hours old bacterial suspensions prepared in NB medium were spread on Petri plates containing solid NA medium. Small filter paper discs immersed in each antibiotic concentration were placed on the media and plates were incubated at 26±2°C for 24 hours. Each treatment was replicated five times and zone of inhibition was measured from each treatment.

## IN-PLANTA ASSAYS

### Experiment 1: Effect of seed bacterization on germination and vigor index in chilli

For this, chilli seeds (variety: Long green) were bacterized by immersing surface sterilized seeds in 24 hours old bacterial inoculum prepared in 25ml LB medium (bacterial concentrations 10^6^, 10^7^ and 10^8^ cfu ml^-1^) by gently shaking on shaker for 2 hours. Ten seeds / Petri plate were placed on two layers of moistened filter paper in each 9-cm plate and were incubated at 28±2 °C in growth room (Seleim et al., 2011). Filter papers were kept moist by adding 5ml autoclaved distilled water when needed. Seeds soaked in only autoclaved distilled water were kept as control. Data on growth parameters and vigor index was recorded after 20 days of incubation. Experiment was performed with five replications for each treatment. Seed germination percentage (GP) and vigor index (VI) were recorded by the formulas: 

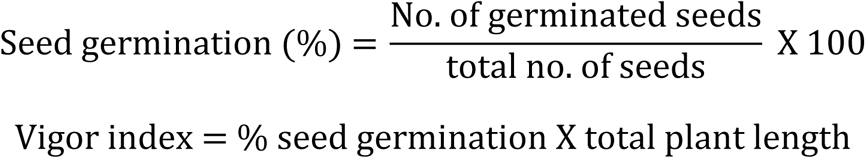

### Experiment 2: Evaluation of bacterial isolates for the induction of defense related enzymes

Bacterial antagonists were evaluated for defense related enzyme induction ability in a pot experiment on chilli (variety: Long green) under natural conditions in a net house. Plastic pots of 1.5 L capacity were filled with autoclaved sandy loam textured soil and flooded with 20ml sporangial suspension of *P. myriotylum* (1×10^3^ sporangia/ml). Fifteen days old healthy seedlings were dipped for 2 hours in overnight bacterial suspension (10^8^ cfu/ml) in LB medium before shifting in pots containing infested soil. Five seedlings per pot were sown in three repeats and placed under net house conditions at 28±2 °C and 80% relative humidity. The experiment was performed with ten treatments viz., T1 (*P. myriotylum* as negative control), T2 (4a2 – *Flavobacterium* spp.), T3 (JHL-8 – *Bacillus megaterium*), T4 (JHL-12 – *P. putida*) T5 (1C2 – *B. cereus*), T6 (RH-24 – *B. subtilus*), T7 (1D – *B. cereus*), T8 (5C – *P. putida*) T9 (RH-87 – *P. libanensis*) T10 (Untreated control) in three repeats. Root tissues were taken at 1, 3 and 5 days intervals after transplant.

### Enzymes extraction and quantification

Representative chilli root samples (2g) were crushed with 4ml of 0.1M sodium phosphate buffer at 4°C in sterilized mortar and pestle. The homogenized solution was centrifuged for 15 minutes at 10,000 rpm and 4°C and supernatant was used for the estimation of PO, PPO, PAL and chitinase activity by spectrophotometry (Anand et al., 2007).

### Peroxidase test (PO)

PO activity was tested by following the methodology adopted by (Hammerschmidt et al., 1982). In this test, peroxidase activity was measured by mixing 0.5ml of enzyme extract with 1.5ml of pyrogallol (0.05M) and 0.5ml of 1% H_2_O_2_ and incubated at room temperature. Absorbance change was noted at 420nm at 30 s interval for 3 minutes against a blank.

### Polyphenol oxidase test

PPO test was performed in accordance with the methodology described by (Mayer et al., 1966). In particular, PPO activity was determined by mixing 200ml of the crude enzyme extract with 1.5ml of 0.1M sodium phosphate buffer, 200ml of 0.01M catechol was added to start the reaction and absorbance was recorded at 495 nm wavelength.

### Phenylalanine ammonialyas test (PAL)

PAL activity test was performed in accordance with (Whetten and Sederoff, 1992). In particular, PAL activity was assessed by mixing 100µl of enzyme, 500µl of 50mM Tris HCL, and 600µl of 1mM L-phenylalanine followed by incubation for 1 hour. Reaction was stopped by adding with 2N HCL followed by adding 1.5ml toluene in the mixture, vortexed for 30 s, and centrifugation at 1000 rpm for 5 minutes. Toluene fraction carrying trans-cinnamic acid was separated. The toluene phase was estimated at 290 nm wavelength against the toluene as blank, and a standard curve was constructed with graded amounts of cinnamic acid in toluene.

### Experiment 3: Testing of bacteria for disease suppression and plant growth promotion (PGP) traits in pot trials

Potential of antagonistic bacteria to suppress the damping-off disease and plant growth promotion effect was tested under natural environmental condition in net house conditions in a repeated experiment. In this study, zoospores of *P. myriotylum* were obtained by following the procedure previously described by (Rahimian and Banihashemi, 1979) and concentration was maintained at 1 x 10^6^ zoospores/ml using hemocytometer. Plastic pots (1.5 L) were filled with sterilized soil/peat (75% : 25% ratio) and 100ml zoospore suspension of *P. myriotylum* was added in soil before sowing the chilli seeds (Variety: Long green). Prior to sowing, surface sterilized chilli seeds were soaked individually for 2 hours in bacterial suspensions (10^8^ cfu/ml) prepared in Luria-Bertani (LB) medium (Sigma-Aldrich, USA). Un-inoculated seeds without *P. myriotylum* and rhizobacteria were kept as untreated control (UTC) while the seeds inoculated with *P. myriotylum* were kept as negative control (NC). All other plant management practices were kept same for all the treatments. Experiments were carried out with five replications for each treatment. Pots were kept under net house conditions and damping-off disease incidence and seed germination percentage was recorded after 15 days of sowing while data on PGP was taken 30 days after sowing the seeds.

### Statistical data analysis

Statistical data analysis was performed using Statistix 8.1 software and MS Excel 2010. All the experiments were performed in completely randomized design (CRD) with replicated treatments. All the experiments were repeated at least two times to confirm the results. Mean values for each treatment were calculated, and all the treatment means were compared via Analysis of variance (ANOVA) test using the least significant differences (LSD) at 5% probability (P≤0.05). Correlation was studied in Microsoft Office Excel 2010.

## RESULTS

### Pathogen inoculum of *P. myriotylum*

A total of 13 isolates of *P. myriotylum* were recovered from infected chilli roots showing the characteristics symptoms of damping-off disease on corn meal agar medium (CMA) and were identified on the basis of morphological characters i.e. coenocytic hyphae bearing lobate sporangia (7 to 15μm wide), knob-like appressorium, vesicles (43 to 52μm in diameter) bearing 29 to 45 zoospores/vesicle, encysted zoospores (10 to 12μm in diameter), terminal oogonia (30 to 38μm in diameter), crooked necked antheridia (4 to 7 antheridia per oogonium), and aplerotic oospores (25 to 31μm in diameter) as described by (Drechsler, 1943) **(Fig. 1)**. Internal transcribed spacer regions (ITS1 and ITS2) were amplified by PCR and final sequences were submitted to GenBank database under the accessions; MF143429 and MF143430. BLAST analysis of approximately 700bp fragments showed 99% sequence identity with already published sequence of *P. myriotylum* (accession HQ643704) (Robideau et al., 2011). Both the isolates PMyr-1 and PMyr-2 produced characteristic damping-off symptoms in pathogenicity tests on chilli. This pathogen was previously published as first report of *P. myriotylum* causing damping-off and root rot in chili from Punjab Pakistan.

**Figure 1.**
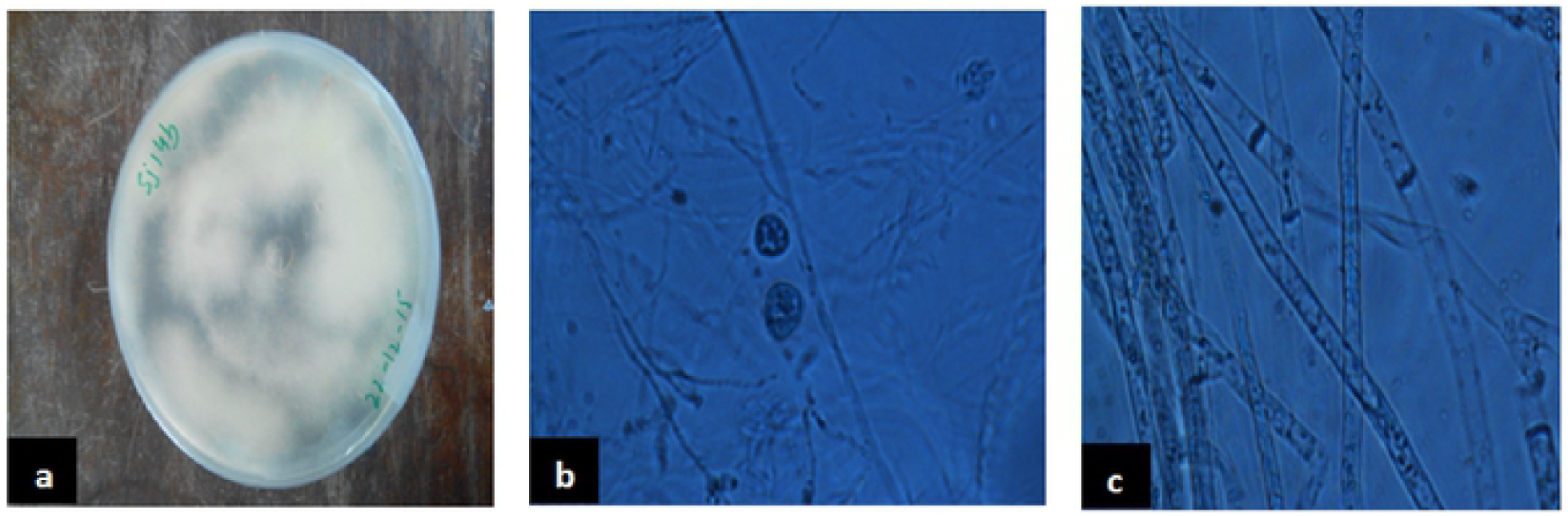
Pure culture of *P. myriotylum* on corn meal agar medium (CMA) supplemented with ampicillin (250 mg/l), rifampicin (10 mg/l), and pimaricin (10 mg/l) (a), sexual fruiting bodies (b), and coenocytic fungal hyphae under microscope.

### Isolation and *in vitro* screening of bacterial isolates against *P. myriotylum*

A total of 110 rhizobacterial isolates were recovered from the healthy chilli roots and rhizospheric soil samples and were screened for antagonistic potential against two highly virulent strains of *P. myriotylum* (PMyr-1 and PMyr-2) isolates in repeated dual culture experiments on PDA medium. Out of all tested isolates, 28 (25.5%) bacterial isolates exhibited varied levels of antagonistic activities (Unpublished data). Data recorded after 48 and 96 hours of incubation showed that out of 28 bacterial isolates, 8 (28.6%) isolates 4a2-*Flavobacterium* spp., JHL-8-*Bacillus megaterium*, JHL-12-*Pseudomonas putida*, 1C2-*B. cereus*, RH-24-*B. subtilis*, 1D-*B. cereus*, 5C-*P. putida* and RH-87-*P. libanensis* exhibited significant antagonistic activity against *P. myriotylum* as compared to control under *in vitro* conditions (**Fig. 2**). Forty-eight hours after inoculation, mycelial growth inhibition percentage in PMyr-1 was ranged from 38.6 to 81.4% (48 hours after inoculation) and 36.1 to 76.7% (96 hours after inoculation), while percentage mycelial growth inhibition in PMyr-2 was 48.5 to 80.6 % and 41.4 to 75.9% after 48 and 96 hours of inoculation respectively. These 8 potential bacterial isolates were further tested by subsequent *in vitro* experiments.

**Figure 2.**
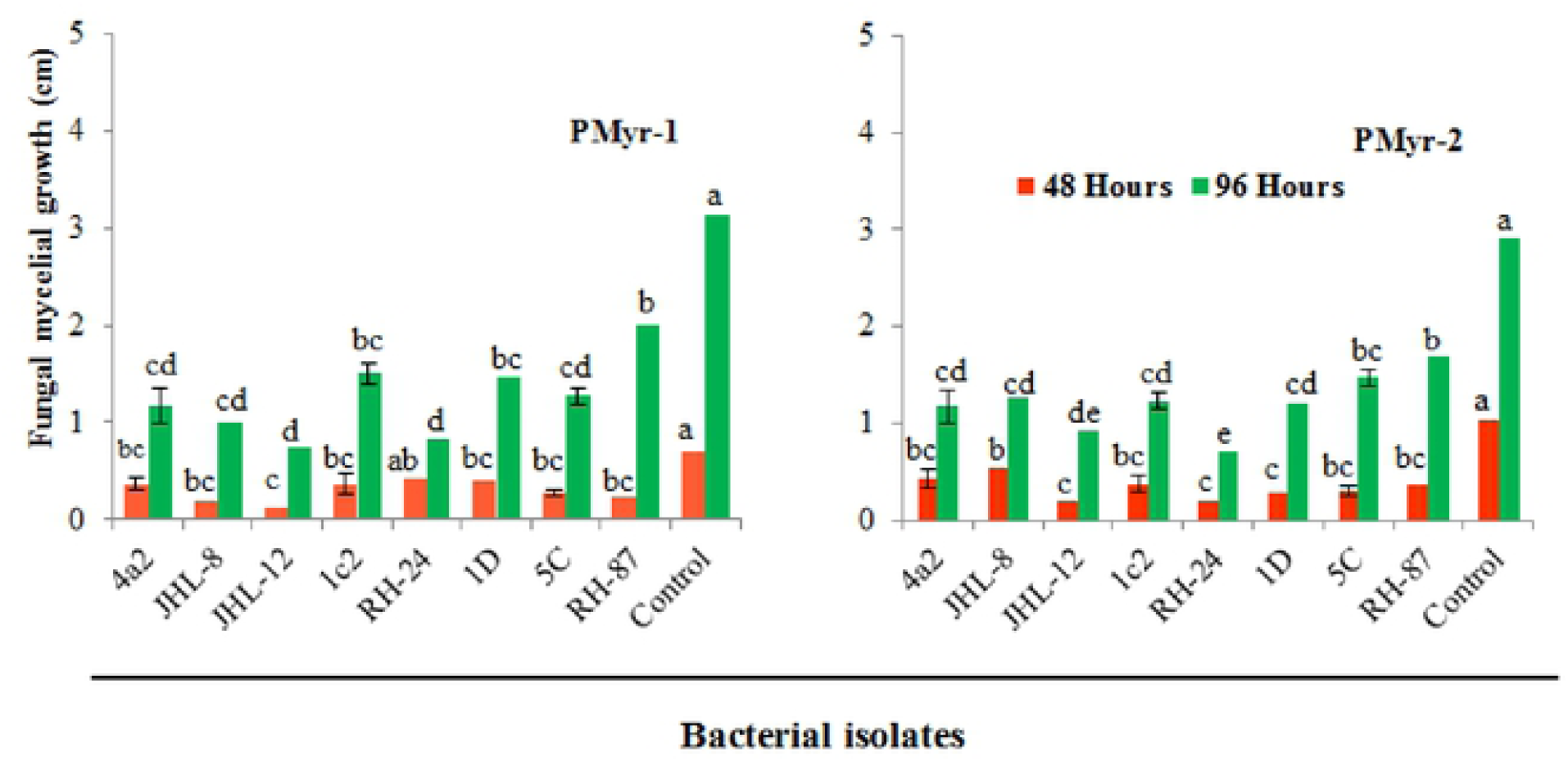
*In vitro* antagonistic activity of rhizobacterial isolates against *P. myriotylum*, the causal agent of damping-off in chilli. Fungal mycelial growth was measured in cm from each treatment. PMyr-1 and PMyr-2 are two virulent strains of *P. myriotylum* while 4a2, JHL-8, JHL-12, 1c2, RH-24, ID, 5C and RH-87 are rhizobacterial isolates. Presented values are the average of three replicates of each treatment. Error bars show standard deviations and letters on each bar represent the significant difference among the values at 5 % level of significance.

### Biochemical featuring of rhizobacterial isolates

The response of all the tested bacterial isolates towards various biochemical tests is presented in **table 1**. Out of 8 bacterial isolates, 4 isolates JHL-8, 1C2, RH-24 and 1D were gram positive while 3 isolates gave fluorescence emission. Four bacterial isolates 4a2, JHL-12, 5C and RH-87 gave positive results for KOH solubility test while all the bacteria isolates were positive for catalase production. In response to levan production test, 2 bacterial isolates, 1D and RH-87 showed positive test results while 3 bacterial antagonists JHL-12, 1C2 and 1D were positive for carbohydrate fermentation reaction. All the tested bacterial isolates showed positive results for oxidase test except RH-87 which was not tested for the response. All the bacterial antagonists except RH-24 exhibited positive response for oxidative fermentative test. Bacterial isolates except 4a2 gave positive results for nitrate reduction and gelatin hydrolysis test. Nitrate reduction and gelatin hydrolysis tests were not performed on RH-87 and 4a2 bacterial isolates respectively.

**Table 1.** Response of rhizobacterial isolates to various biochemical tests.

| Bacterial isolates | GS | FE | KOH | CAT | LP | CF | OXD | OFR | NR | GH |
| --- | --- | --- | --- | --- | --- | --- | --- | --- | --- | --- |
| 4a2 | - | - | + | + | - | - | + | + | - | NA |
| JHL-8 | + | - | - | + | - | - | + | + | + | + |
| JHL-12 | - | + | + | + | - | + | + | + | + | + |
| 1c2 | + | - | - | + | - | + | + | + | + | + |
| RH-24 | + | - | - | + | - | - | + | - | + | + |
| 1D | + | - | - | + | + | + | + | + | + | + |
| 5C | - | + | + | + | - | - | + | + | + | + |
| RH-87 | - | + | + | + | + | - | NA | + | NA | + |

\*Each test result was confirmed in three replications for each bacterial isolate. (+) positive reaction (-) negative reaction and (NA) not tested results for: Gram staining (GS), Fluorescence emission (FE), Potassium hydroxide (KOH), Catalase test (CAT), Levan production (LP), Carbohydrate fermentation (CF), Oxidase test (OXD), Oxidative fermentative (OFR), Nitrate reduction (NR), Gelatin hydrolysis (GH) test.

### Molecular characterization of rhizobacterial isolates

Phylogenetic analysis from 16S rRNA sequences (≈ 1500 bp) of 8 rhizobacterial isolates showed that all the tested bacteria were belonged to *Bacillus, Pseudomonas*, and *Flavobacterium* spp. (**Figure 3**). Bacterial isolates ID and 1C2 had 97 to 98% sequence homology with *Bacillus cereus* (accessions MK606105 and MK648339 respectively) and the sequences of ID and 1C2 were submitted to GenBank database under accession numbers MH393211 and MH393210.1 respectively. The 16S rRNA gene sequence of the bacterial isolate JHL-8 (accession MH393209) showed 99% sequence similarity with *B. megaterium* (accession MG430236). The sequence of RH-24 (accession MH393208) had 99% sequence homology with *B. subtilis* (accession KY000519). Isolate RH-87 (accession MT421780) was closely related to *Pseudomonas libanensis* 99% identity with GenBank accession number DQ095905. Bacterial isolates 5C and JHL-12 had 99% sequence homology with *P. putida* (accessions KY982927 and MF276642 respectively) and the sequences of 5C and JHL-12 were deposited to GenBank database under accession numbers MH371201 and MH371200 respectively. Bacterial isolate 4a2 (accession MT421823) displayed 99% identity with the GenBank sequence of *Flavobacterium* spp. (accession HM745136). The accession numbers of all the tested bacterial antagonists and sequence homology percentage with their reference isolates are given in **Table. 2**.

**Table 2.**
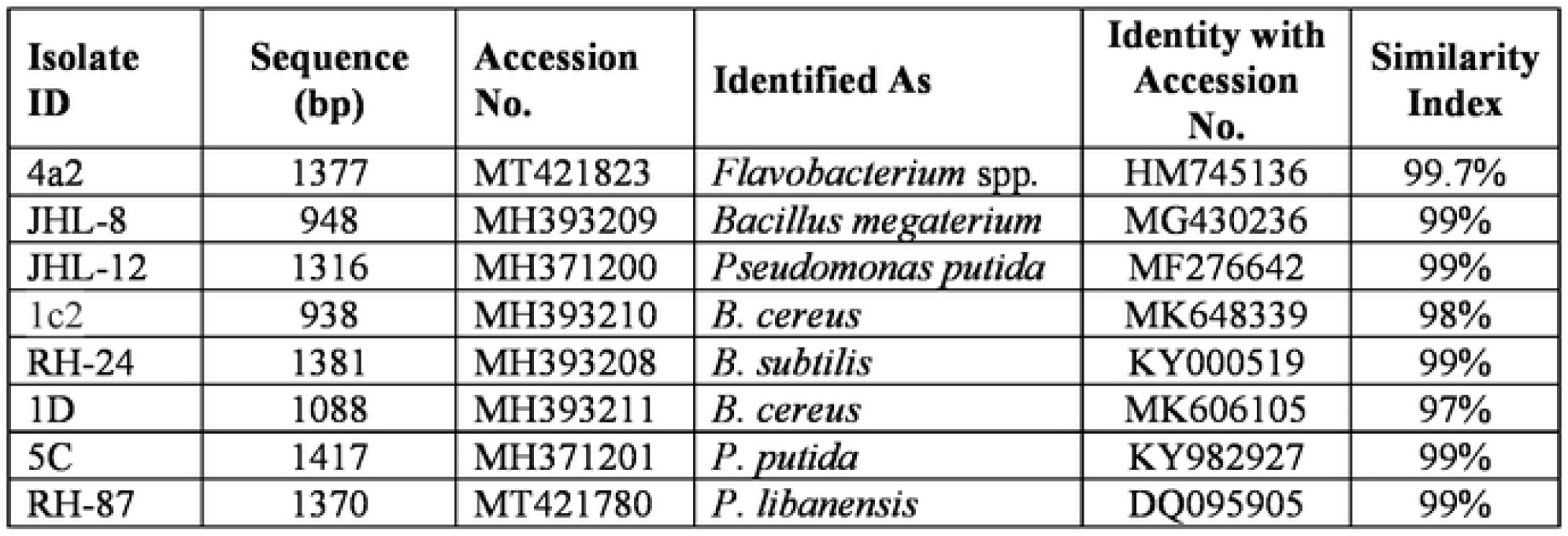
Sequence analysis of 16S rRNA from rhizobacterial isolates and their homology with the reference bacterial.

**Figure 3.**
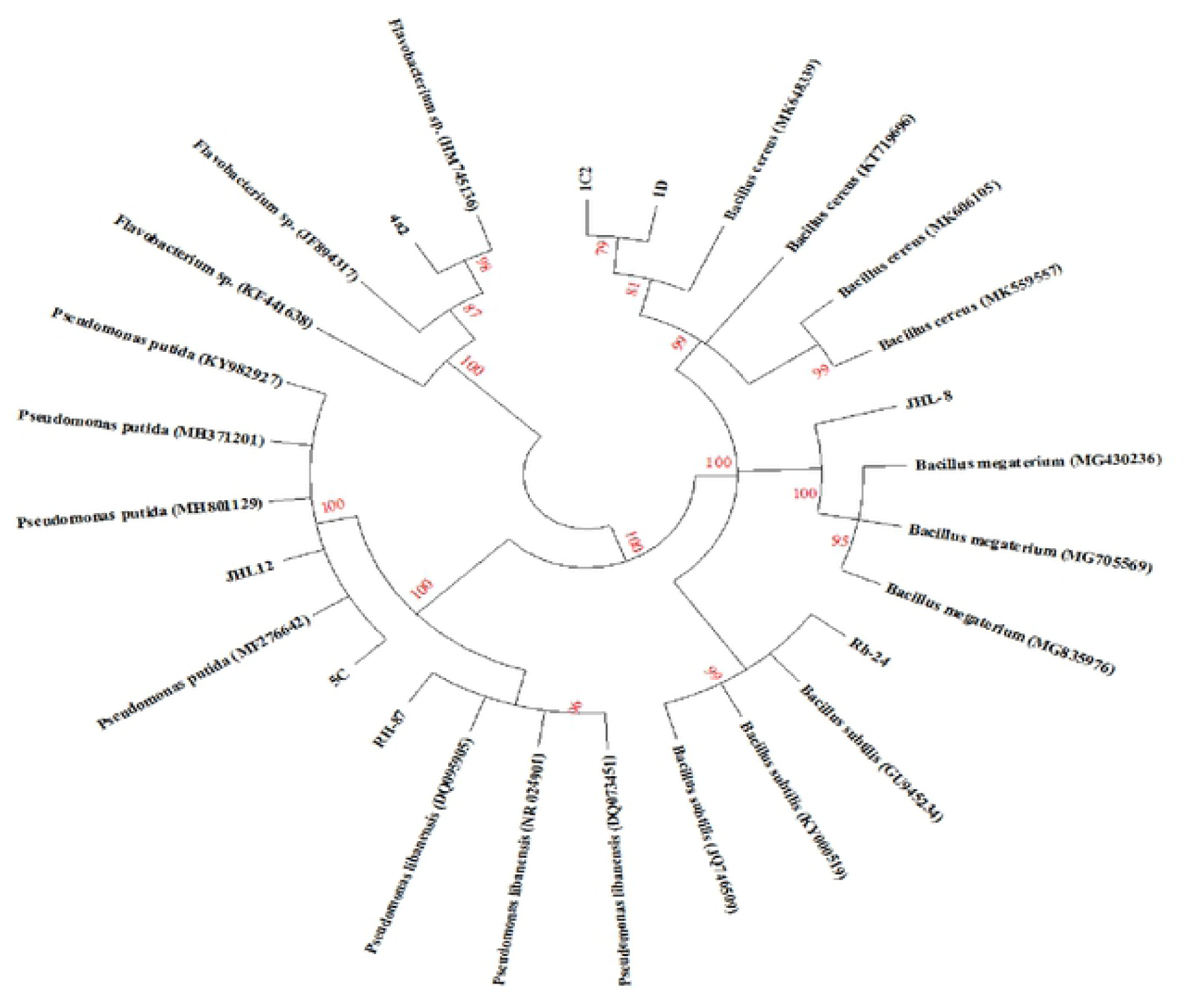
Phylogenetic tree based on the 16S rRNA sequences (≈ 1500 bp) showing the relationships between the representative rhizobacterial isolates and closely associated neighbors. 16s rRNA gene fragments were amplified by PCR protocols using 27F and 1492R primers and amplified products were confirmed by Gel electrophoresis against 1kb ladder. Gel purified DNA products were sent for sequencing and obtained sequences were joined by using DNASTAR software. BLAST analysis was performed to retrieve closely associated bacterial sequences. All the Sequences were aligned in CLUSTAL W program and phylogenetic tree was constructed by using Kumara 2-parameter model (K2 + G) using 1000 replicates as bootstrap values and >70% are labeled.

### Characterization of rhizobacterial isolates for biocontrol and plant growth promotion (PGP) traits

All the rhizobacterial isolates were tested for biocontrol and plant growth promotion (PGP) traits and their responses are given in **table 3**. All the bacterial isolates displayed positive test results for ammonia production as indicated by the brown to yellow color development. All the bacterial isolates were able to hydrolase starch while test was not performed for bacterial isolate 4a2. All the tested bacterial isolates except 1D exhibited positive test results for hydrogen cyanide (HCN) production, and the said test was not performed for 4a2. A halo zone formation around the bacterial growth on Pikovskaya’s agar medium after 96 hours indicated the positive test results for phosphate solublization by the bacterial isolates. *Bacillus* spp. JHL-8 showed highest P-solubilization (103 μgml^−1^) followed by RH-24 (97 μgml^−1^). In case of *Pseudomonas* spp., 5C exhibited maximum P-solubilization (84 μgml^−1^) followed by RH-87 (79 μgml^−1^) and JHL-12 (75 μgml^−1^). *Flavobacterium* spp. Bacterial isolate 4a2 showed maximum P-solubilization (86 μgml^−1^) and the test was not done for bacterial isolate 1D. All the bacterial isolates showed pink-red color development as an indication of IAA production. The spectrophotometry study confirmed the production of IAA by bacterial isolates between 39 μgml^−1^ and 13.4 μgml^−1^. Maximum IAA (39 μgml^−1^) was produced by RH-24 followed by 4a2 (34.1μg ml^−1^) and JHL-8 and 5C (26.7 μgml^−1^) while minimum amount of IAA (13.4 μgml^−1^) was produced by RH-87 respectively. Siderophore production was indicated by the change in color from blue to orange on chrome azurol S agar medium. Among the *Bacillus* spp. RH-24 showed highest siderophores production (27.7%) followed by JHL-8 (23.7%) and 1C2 (23.3%). *Pseudomonas* spp. RH-87 showed highest siderophores production (23.6%) and 5C (19%) while *Flavobacterium* spp. 4a2 showed 18.5% siderophores production.

**Table 3.** Characterization of rhizobacterial isolates for plant growth promoting (PGP) traits.

| **Bacterial isolates** | **AP** | **SH** | **HCN** | **P-Solubilization ($\mu\text{g ml}^{-1}$)** | **IAA production ($\mu\text{g ml}^{-1}$)** | **SDP (%)** |
| --- | --- | --- | --- | --- | --- | --- |
| **4a2** | + | ND | ND | $86 \pm 4.33^b$ | $34.1 \pm 1.7^a$ | $18.5 \pm 1.1^d$ |
| **JHL-8** | + | + | + | $103 \pm 3.53^a$ | $26.7 \pm 2.3^b$ | $23.7 \pm 2.0^{ab}$ |
| **JHL-12** | + | + | + | $75 \pm 2.89^c$ | $19.6 \pm 1.4^c$ | $17.3 \pm 1.4^d$ |
| **1c2** | + | + | + | $81 \pm 3.28^{bc}$ | $13.9 \pm 1.7^{cd}$ | $23.3 \pm 0.8^{bc}$ |
| **RH-24** | + | + | + | $97 \pm 4.36^a$ | $39.0 \pm 1.5^a$ | $27.7 \pm 1.5^a$ |
| **1D** | + | + | - | ND | $19.3 \pm 2.8^{cd}$ | $17.1 \pm 0.9^d$ |
| **5C** | + | + | + | $84 \pm 2.31^{bc}$ | $26.7 \pm 2.2^b$ | $19.0 \pm 1.7^{cd}$ |
| **RH-87** | + | + | + | $79 \pm 3.93^{bc}$ | $13.4 \pm 1.5^d$ | $23.6 \pm 1.6^{ab}$ |
| **LSD** |  |  |  | **10.062** | **5.8787** | **4.2705** |

(+) positive (-) negative test, AP (Ammonia production), SH (Starch hydrolase), HCN (Hydrogen cyanide production), PHS (Phosphate solubilization), IAA (Indole acetic Acid production), SDP (Siderophore production %). Data are average values of three replicates for each bacterial isolate. ( ± ) standard error values.

### Multiple antibiotic resistance of rhizobacterial isolates

All the bacteria displayed varying level of resistance and susceptibility against all tested antibiotics (**Fig. 4**) and zone of inhibition around the bacterial cultures were measured (**Fig. 5a to 5e**). All the rhizobacterial isolates showed no tolerance against Streptomyces at all dose levels except *Bacillus subtilis* (RH-24) which showed resistance up to 400ppm dose level. Against Ampicillin, all the tested bacteria showed resistance to all the dose levels except 1C2 and RH-87 which showed little susceptibility at two highest dose levels. When tested against Penicillin G, bacterial isolates, RH-24, 5C, 1C2, 4a2, JHL-8 and JHL-12 were found resistance at all the dose levels while 1D and RH-87 showed little susceptibility at 500ppm dose level. Most of the tested bacteria showed no resistance against Rifampicin at all six dose levels however, RH-24 showed resistance against Rifampicin up to 300ppm dose level and 4a2 showed little susceptibility against Rifampicin at 500ppm. Bacterial isolate RH-24 showed highest tolerance against Vancomycin at all the dose level while all other tested bacterial isolates showed varied levels of susceptibility response at different dost levels. The tested isolates ID, 1C2 and RH-87 showed resistance response up to 300ppm dose level of Vancomycin while the bacterial isolates 4a2, JHL-8 and JHL-12 showed maximum susceptibility.

**Figure 4.**
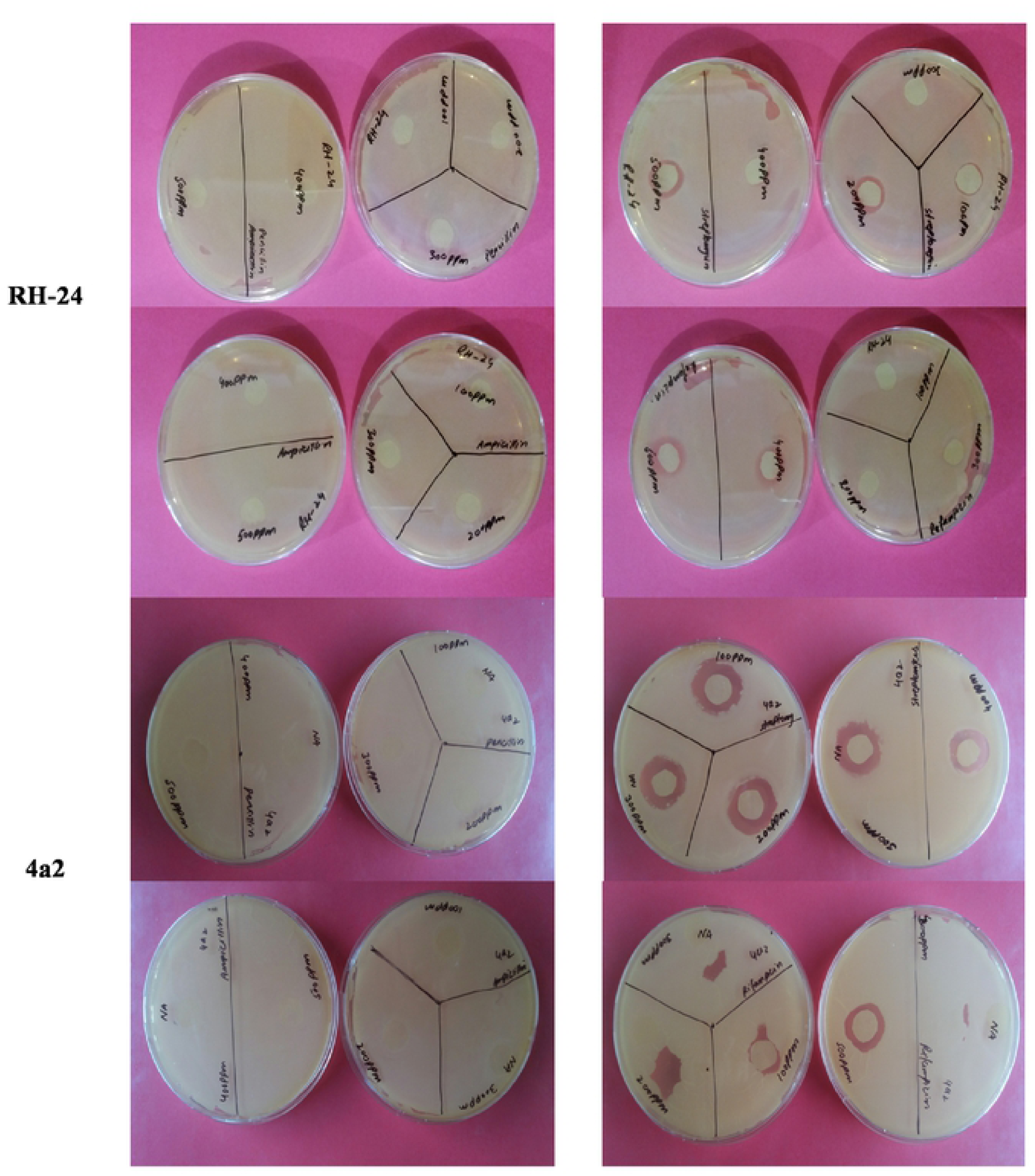
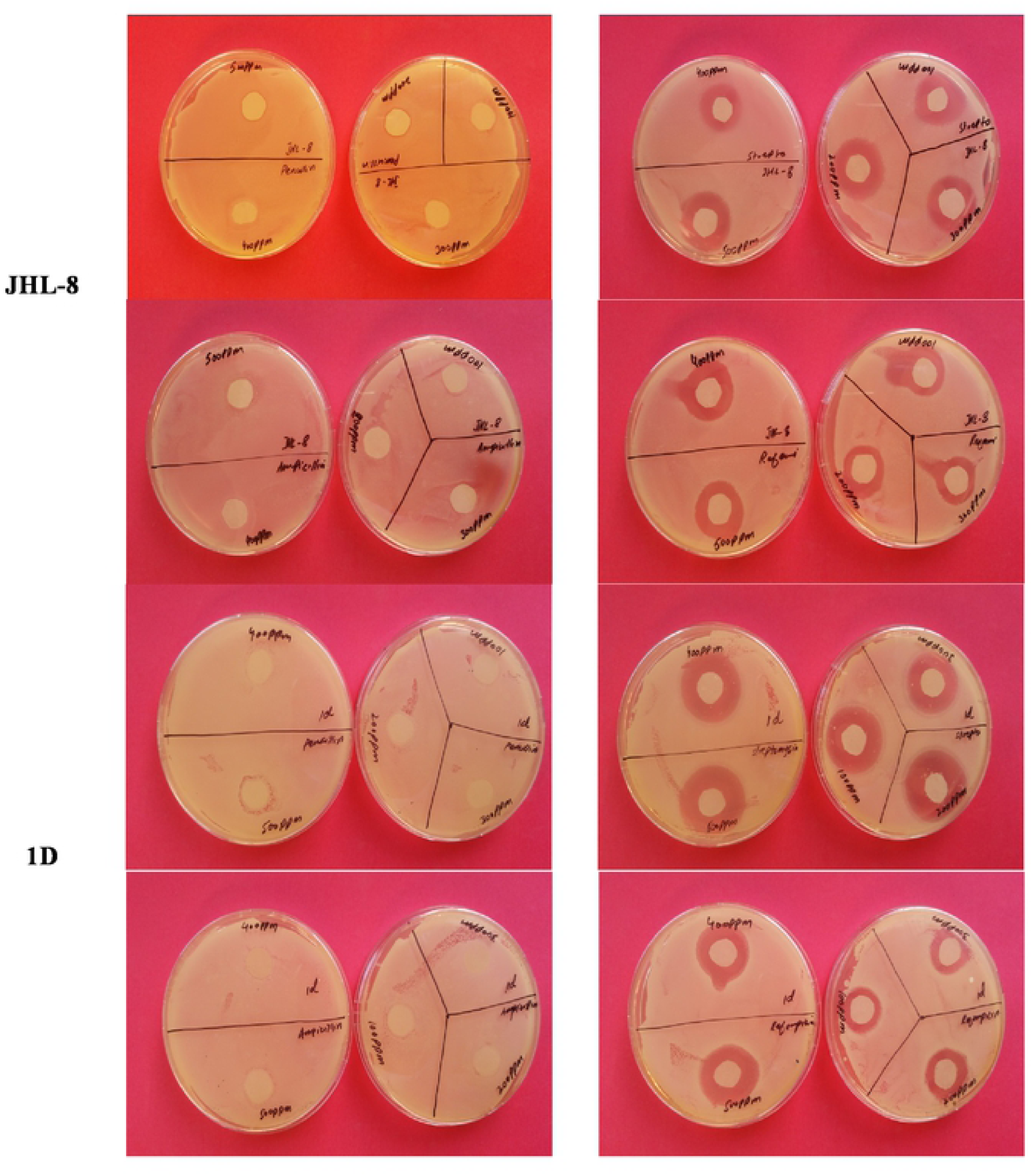
Multiple antibiotic resistance assays to assess the bacterial resistance and susceptibility levels against Streptomyces, Ampicillin, Rifampicin, Penicillin G and Vancomycin at 0ppm, 100ppm, 200ppm, 300ppm, 400ppm and 500ppm concentrations. Zone of inhibition (cm) was measured from each replicated plate 24 hours after incubation. Rh-24, 4a2, JHL-8 and 1D are the tested bacterial isolates.

**Figure 5.**
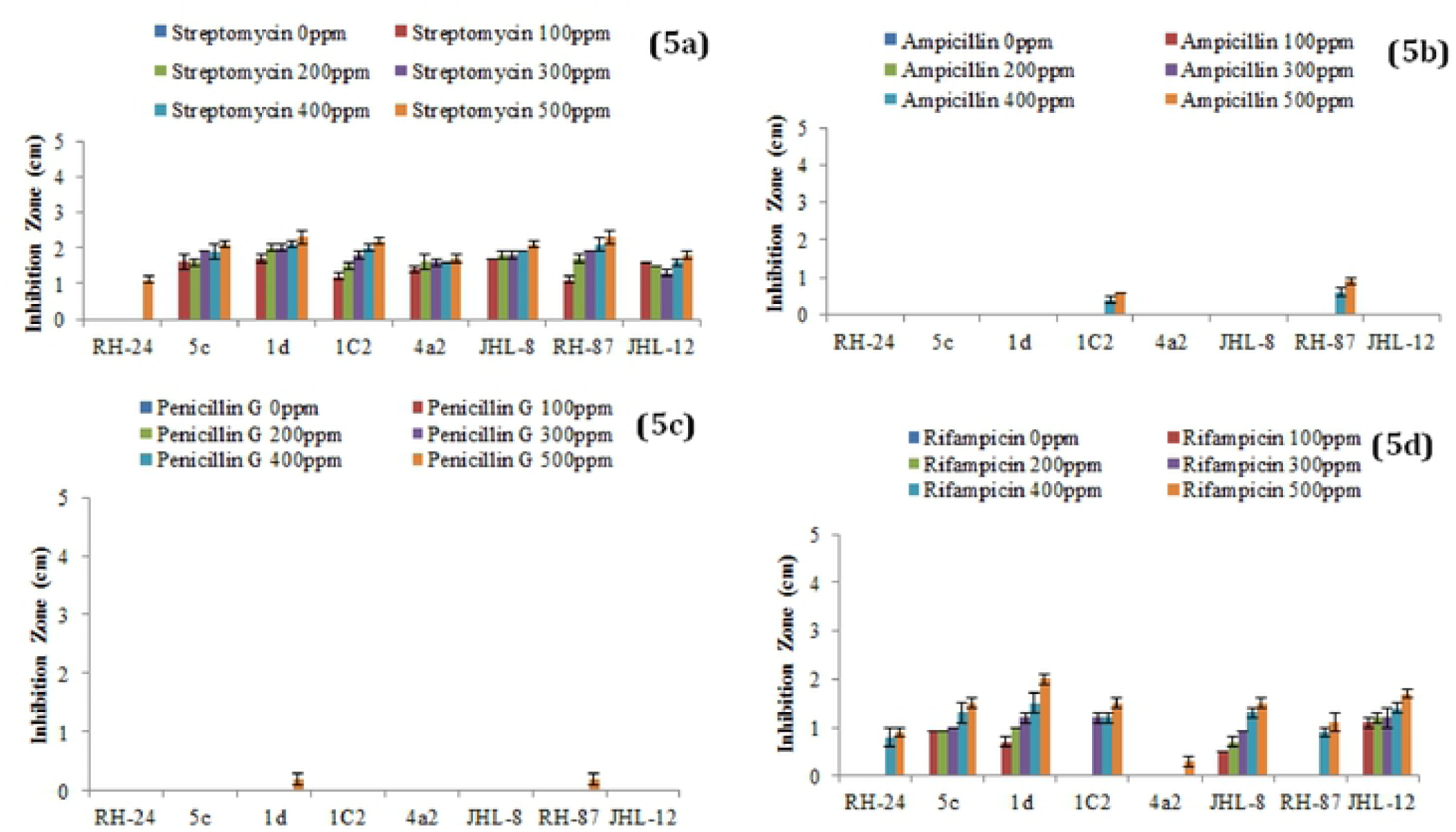
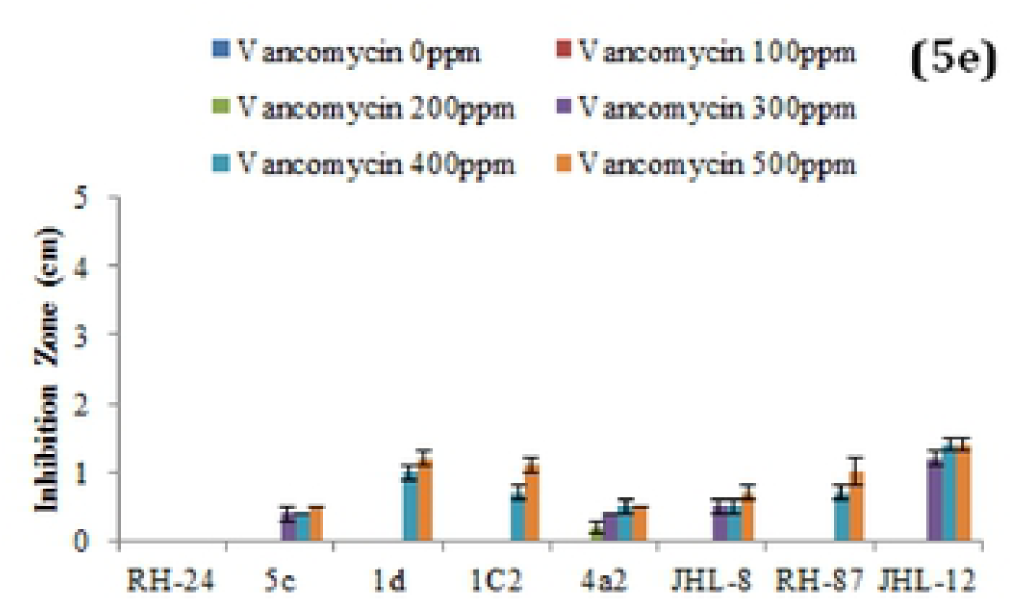
Multiple antibiotic resistance assays to test the bacterial resistance and susceptibility levels against Streptomyces, Ampicillin, Rifampicin, Penicillin G and Vancomycin at six different concentration levels. Each treatment was tested in five replications and bar graphs are made by using average values for each treatment. Error bars on each bar represent the standard error (SE). Rh-24, 5C, 1D, 1C2, 4a2, JHL-8, RH-87 and JHL-12 are the tested bacterial isolates.

## IN-PLANTA ASSAYS

### Effect of bacterial inoculants on chilli seed germination

The effect of bacterial seed treatment upon seed germination and plant growth parameters (PGP) varied with different bacterial isolates (**Fig. 6a–6d**). All the bacterial isolates produced significant effects on seed germination percentage and PGP compared to control treatment. None of the tested bacterial isolate at any applied concentration level showed reduction in seedling germination and phytotoxic effects on chilli seedlings. Maximum plumule length was recorded 11.8cm in chilli seedling inoculated with 1D at 10^7^ *cfu* followed by RH-24 (9.6cm) over control. Maximum radical length was recorded 4cm for 5C at 10^8^ *cfu* followed by 1C2 and 1D while minimum radical length was recorded for RH-87 at all the tested concentrations compared to control. Vigor index was significantly increased in bacterized chilli seed over untreated control.

**Figure 6.**
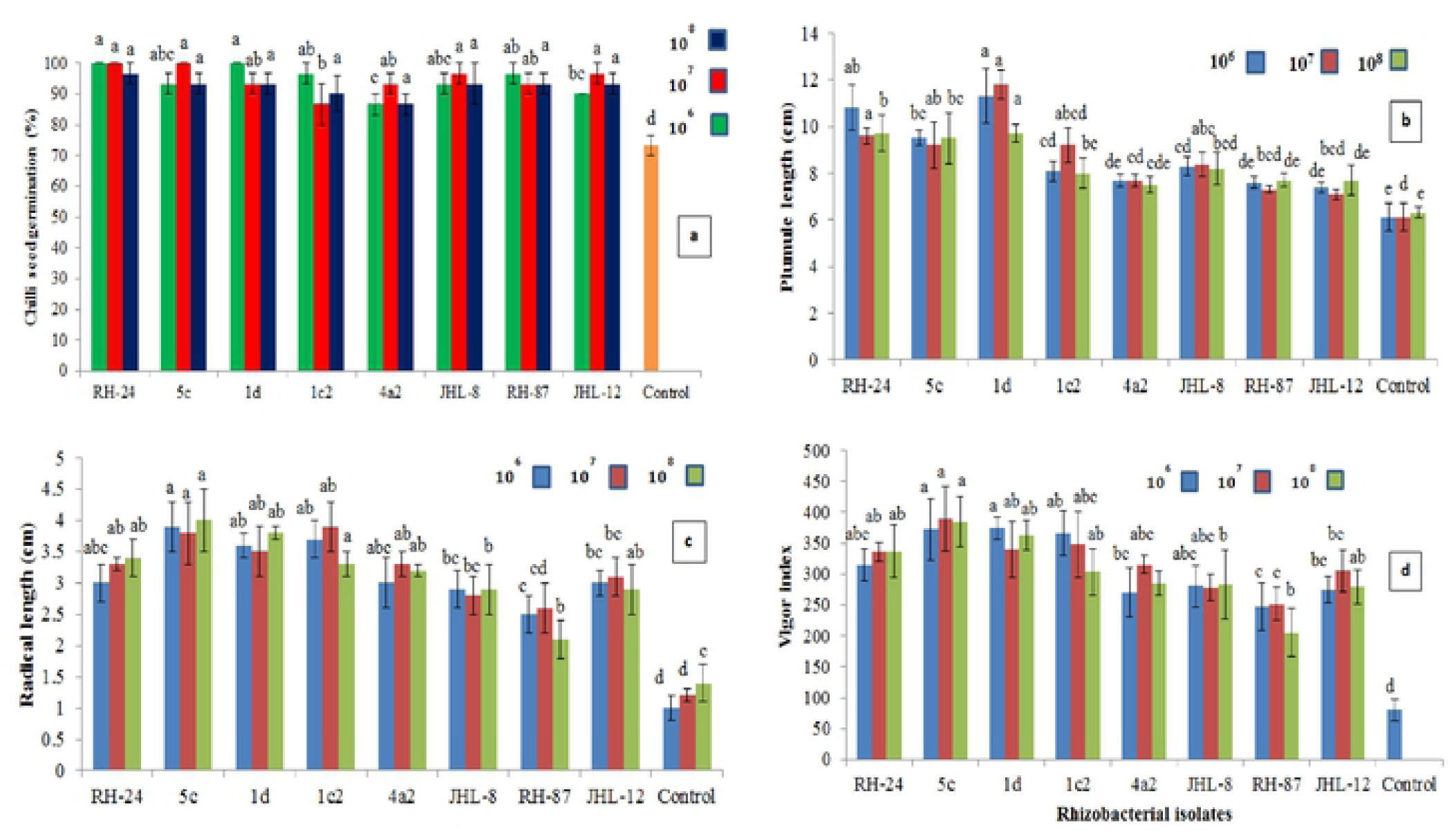
Effect of bacterial inoculants on chilli seeds **(a)** seed germination percentage, **(b)** plumule length, **(c)** radical length, **(d)** vigor index. Chilli seeds were treated with bacterial strains; *Flavobacterium* spp. 4a2, Pseudomonas spp. JHL-12, 5C, RH-87, and *Bacillus* spp. JHL-8, 1C2, RH-24, 1D while the seeds in control treatment were dipped in double sterilized water only. Seeds were grown on moist Whatman filter paper No. 41 in sterile Petri plates under controlled conditions. Each treatment was replicated three times and bars are made from the average of three values. Error bars on each bar represent the standard error (SE).

### Induction of defense related enzymes in chilli plants

All the tested bacterial isolates significantly induced defence related enzymes in chilli seedlings under the pathogen presence **(Table 4)**. In chilli seedlings treated with rhizobacterial suspensions, an increase in Peroxidase (PO) activity was observed 3 and 5 days after inoculation (DAI). Maximum PO activities were recorded in seedlings treated with *Flavobacterium* spp. 4a2 followed by *Bacillus* spp. JHL-8 and RH-24 and PO activates were recorded almost three folds higher than negative control (NC) and untreated control (UC). The increased activity of Polyphenol oxidase (PPO) was observed on 5^th^ day after inoculation in seedlings treated with *Bacillus* spp. RH-24 followed by *Flavobacterium* spp. 4a2 and *Pseudomonas* spp. 5C as compared to negative and untreated control. Phenylalanine ammonia-lyase (PAL) activities were observed high in all the chilli seedlings treated with bacterial isolates. Maximum PAL activates were observed in seedlings inoculated with *Bacillus* spp. RH-24 followed by JHL-8 and ID. Phenylalanine ammonia-lyase (PAL) activities were recorded almost 3 folds high in all the bacterial treated seedlings as compared to negative and untreated control.

**Table 4.**
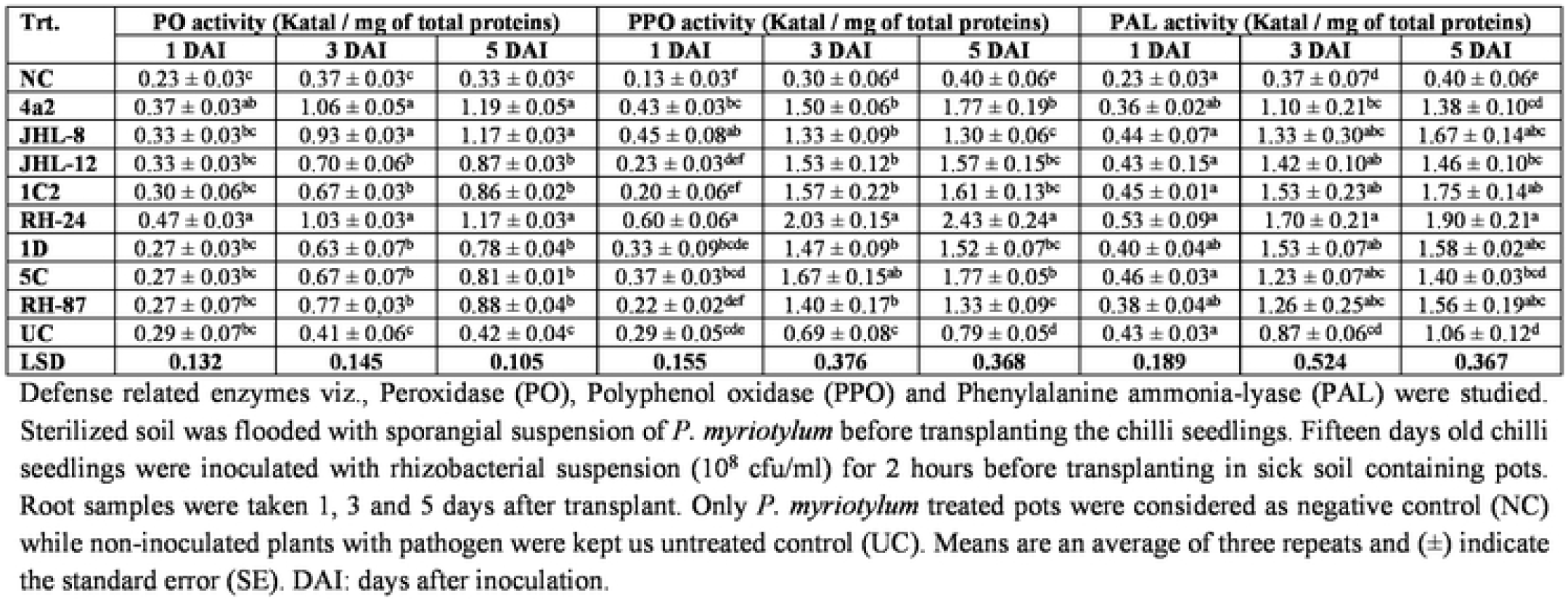
Effect of bacterial isolates on defense related enzymes induction in chilli seedlings inoculated with antagonistic rhhizobacterial isolates.

### Testing of rhizobacterial isolates for disease suppression and plant growth promotion traits in pot trials

Keeping in consideration the biocontrol and plant growth promotion traits, rhizobacterial isolates were screened for disease suppression and plant growth promotion in chilli plants under open environment and data on disease suppression, seed germination and plant growth traits was recorded. All the tested bacterial isolates significantly improved seed germination in the presence of pathogenic fungi in pot soil. Maximum seed germination of 96% was produced by *Bacillus subtilis* – RH-24 followed by *Flavobacterium* spp. – 4a2 (91%) and *B. cereus* – 1D (89%) while *B. megaterium* – JHL-8 showed least effect on seed germination (70%) over negative control – NC (47%). All the tested bacterial isolates suppressed the pathogenic fungi and significantly lower the seedling mortality ranging between 4.4–31% as compared to negative control where seedling mortality was recorded 53% (**Fig. 7a**). A significant increase in shoot length (P≤0.05) was seen in all the treatments of bacterial isolates. Significant increase in shoot length (24.4 cm/plant) was observed in pots treated with *Bacillus subtilis* – RH-24 followed by *Flavobacterium* spp. – 4a2 (18.4 cm/plant) over untreated control treatment-UTC (11.1 cm/plant) and negative control-NC (5 cm/plant). All the bacterial antagonists significantly enhanced the root length ranging 5.6cm – 7.4 cm/plant over untreated control - UTC (5 cm/plant) and negative control - NC where the root length was recorded 2.6 cm/plant (**Fig. 7b**). An increase in fresh shoot weight (P≤0.05) was observed in all the pots treated with bacteria isolates and increase in fresh shoot weight was ranging 1.8g–3.0g as compared to UTC (1.6g) and NC (1.1g). Data recorded from all the bacterial treated pots showed that the increase in the fresh root weight was ranging 0.96g–2.3g as compared to un-inoculated pots (0.93g) and negative control (0.4g) as given (**Fig. 7c**). A significant increase in dry shoot weight (P≤0.05) was also ranged from 1.1g– 2.1g over un-inoculated pots (0.93g) and negative control (0.4g). Among all the tested isolates *Bacillus subtilis* – RH-24 showed highest increase in dry shoot weight (2.1g) while *P. libanensis* – RH-87 showed the least significant increase in dry shoot weight (1.1g). Data on dry root weight displayed that all the tested bacteria isolates had significantly increased the dry root weight (P≤0.05) in chilli plants ranging 0.36g–1.13g over untreated control (0.46g) and negative control (0.16g) (**Fig. 7d**). A positive correlation was observed between shoot, root length and dry shoot, root weight **(Fig. 7e and 7f)** whereas negative correlation was recorded between dry shoot, root weight and chilli seedling mortality percentage **(Fig. 7g and 7h)**.

**Figure 7.**
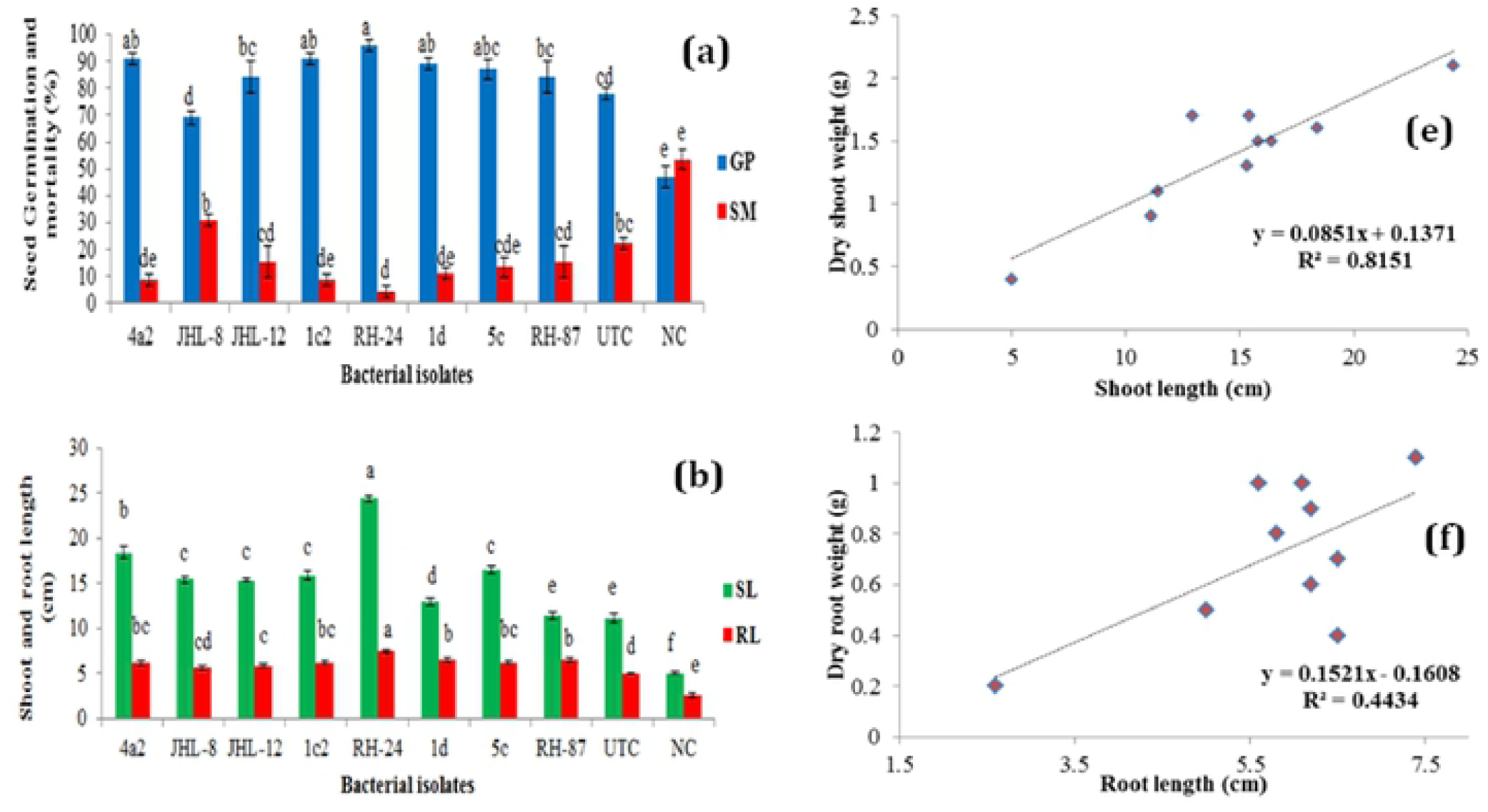
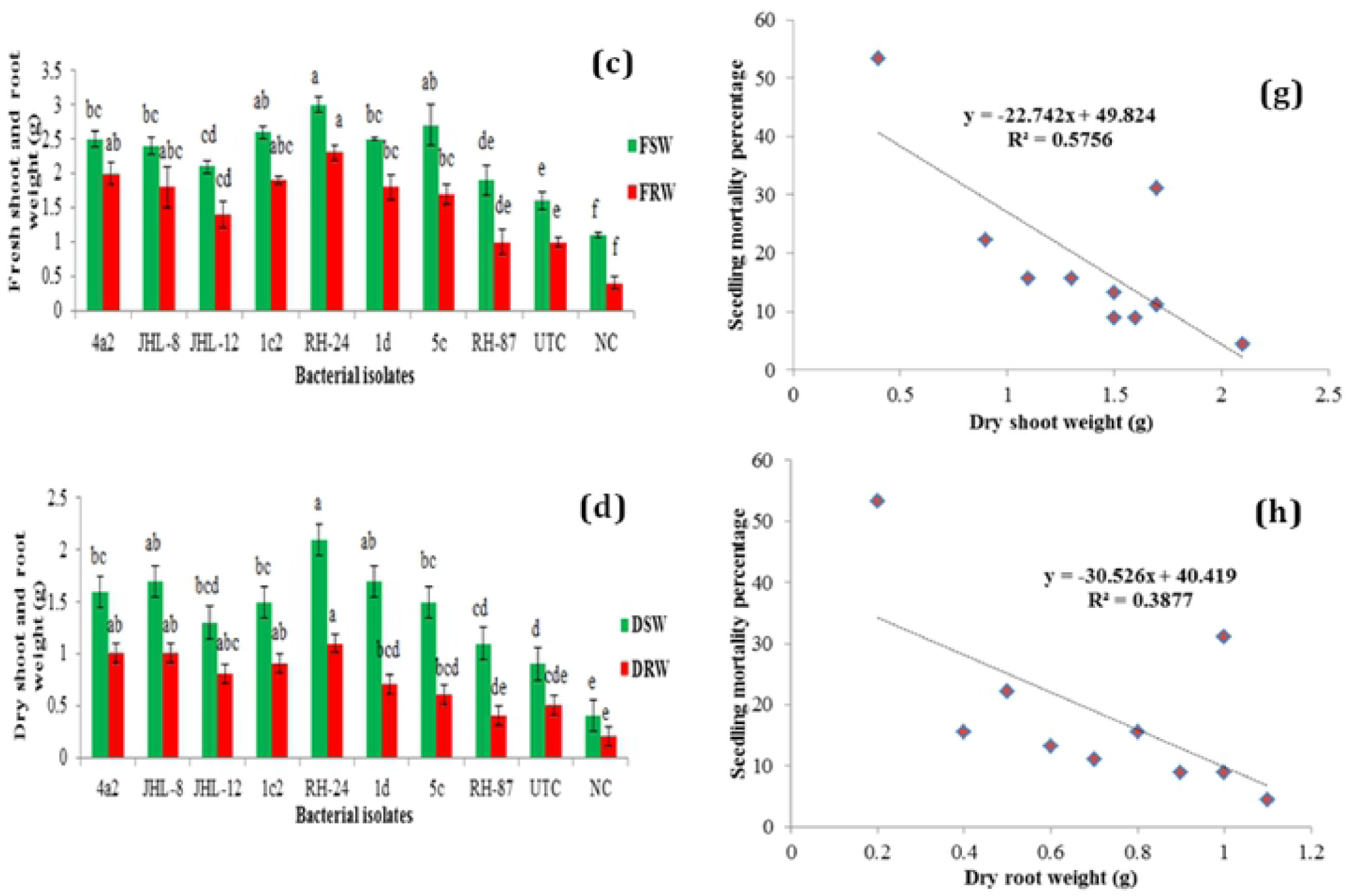
Evaluation of rhizobacterial isolates for the suppression of *Pythium myriotylum* and plant growth promotion effects on chilli seeds in pot trials. Zoospores of *P. myriotylum* were added into the soil and bacterized chilli seedlings were sown in sick soil containing pots in three repeats. Error bars show standard deviations and letters on each bar represent the significant difference among the values at 5 % level of significance. **(a)** seed germination and seed mortality percentage, **(b)** shoot and root length, **(c)** fresh shoot and root weight, **(d)** dry shoot and root weight, **(e)** positive correlation between shoot length and dry shoot weight, **(f)** positive correlation between root length and dry root weight, **(g)** negative correlation between dry shoot weight and seedling mortality percentage, **(h)** negative correlation between dry root weight and seedling mortality percentage.

## DISCUSSION

Chilli pepper is an influential cash crop cultivated across the world, particularly in Asian countries. It belongs to Solanaceae family which contains more than 3000 plant species. Chilli covers 20% of the total vegetable producing area in Pakistan and is majorly cultivated in Sindh province, Pakistan. It is eaten as fresh, dry or processed spices and its hot taste is due to the presence of a compound capsaicinoid. Chilli fruits are cholesterol free and also a serve as a source of folic acid, potassium, vitamins A, B, C, phenolics and carotenoids. Chilli crop is vulnerable to the attack of various diseases at various growth stages. Damping-off by *Pythium* spp. cause decay of germinating seeds and growing seedlings at pre and post-emergence growth stages, and represents huge yield constraints both in nurseries and fields (Erwin and Ribeiro, 1996) and estimated yield loses could be ranging from 5 to 80% under favorable conditions (Lamichhane et al., 2017).

During a field survey in major chilli growing areas in Punjab, Pakistan, from September 2015 to November 2016, young symptomatic chilli seedlings (15 to 30 years old) showing the damping-off, reduced growth, wilting, water soaked lesions, brown discoloration, and root rot (Horst, 2013) were collected. A total of 30 infected root samples were processed for the isolation of *Pythium* spp. onto corn meal agar (CMA) medium amended with ampicillin (250 mg/l), rifampicin (10 mg/l), and pimaricin (10 mg/l) (Jeffers and Martin, 1986), and incubated at 28±2°C (Koehler et al., 2017) for 7 days, and a total of 13 *Pythium* isolates were purified. The morphological characters; coenocytic hyphae with lobate sporangia, knob-like appressorium, zoospores diameter, oogonia diameter, antheridia, and aplerotic oospores characters of the isolates fit well with descriptions of *Pythium myriotylum* Drechsler (Drechsler, 1943).

For pathogenicity assay, seven days old cultures of *Pythium myriotylum* grown on CMA medium were processed for zoospores production by following the methodology previously described by (Rahimian and Banihashemi, 1979). Assay was performed under greenhouse conditions and disease data was recorded fifteen days after sowing the chilli seeds. The inoculated plants produced symptoms of damping-off and root rot, whereas control plants remained asymptomatic. Virulence of *P. myriotylum* on has been reported in various studies (Kageyama and Ui, 1983; Okada, 2003; Tomioka et al., 2013). Two most aggressive isolates PMyr-1 and PMyr-2 from the pathogenicity assay were subjected to molecular characterization. The internal transcribed spacer regions ITS1 and ITS2 were amplified by RCR as described by (White et al., 1990). The amplified sequences (approximately 700-bp) were submitted to the GenBank nucleotide database under the accession numbers MF143429 and MF143430. Sequences of both the isolates showed 99% sequence homology with HQ643704 accession of *P. myriotylum* (Robideau et al., 2011). Similar results were reported by other studies (Tomioka et al., 2013).

Among various disease management practices, chemical seed treatment is extensively adopted strategy as reported (Kandel et al., 2016; Rothrock et al., 2012), and variety of chemicals have been used as seed dressers to remove the pathogens from seeds (Mancini and Romanazzi, 2014). It is an effective practice but can cause adverse effects on seed germination and can cause phytotoxicity (du Toit, 2004). Non-judicial application of these synthetic chemicals is a potential threat to human and environment (Ouyang et al., 2016), noxious to the beneficial rhizosphere microbes (Hussain et al., 2009), pest resistance development (Onstad, 2013) and increasing public security in many countries (Bourguet and Guillemaud, 2016). Many of these chemicals are declared as carcinogen in many countries (Bressa et al., 1997). Keeping in view above mentioned facts and to minimize the dependency on synthetic agrochemicals, scientists have devised alternative, eco-friendly approaches to manage the crop diseases in a more sustainable way.

Of the integrated strategies, the use of PGPR in disease suppression and plant growth promotion (PGP) is widely adopted strategy. Utilization of PGPR in disease suppression and plant growth promotion has been reported on various crops viz., wheat (Abbasi et al., 2011), rice (Yasmin et al., 2016), okra (Begum et al., 2012), cucumber (Islam et al., 2016), sweet pepper (Sid et al., 2003), red pepper (Lim, 2010), avocado (Cazorla et al., 2007), potato (Kenawy et al., 2019) and tomato (Szczech and Shoda, 2004). Rhizobacteria belonging to *Pseudomonas, Azospirillum, Azotobacter, Klebsiella, Enterobacter, Alcaligenes, Arthobacter, Burkholderia, Bacillus*, and *Serratia* spp. enhance plant growth (Souza et al., 2015) and are used as bio-control (El-Sayed et al., 2014; Labuschagne et al., 2010).

Present study was carried out to screen potential antagonistic PRPG isolated from the chilli rhizosphere for the biocontrol of damping-off and plant growth promotion in chilli. Biological control of *Pythium* spp. can be due to antagonistic ability of the rhizobacteri (Clark, 2006; Elazzazy et al., 2012). Out of 110 rhizobacterial isolates, 28 (22.7%) isolates showed varied level of antagonistic potential and out of these 28 bacterial isolates, 8 (28.6%) isolates 4a2, JHL-8, JHL-12, 1C2, RH-24, 1D, 5C and RH-87 showed significantly high antagonistic potential against two highly virulent strains of *P. myriotylum* PMyr-1 and PMyr-2. Antagonistic potential of tested bacterial isolates could be due to antibiosis as various antibiotics have been previously identified and reported (Nielsen and Sørensen, 2003; Raaijmakers et al., 2002). Bacterial isolates showing highest antagonistic potential were subjected to biochemical featuring. In this study, out of 8 bacterial isolates, 4 isolates were gram positive and others were gram negative.

Bacterial isolates belonging to *Bacillus* and *Flavobacterium* spp. showed negative results for fluorescence emission test, while those belonging to *Pseudomonas* spp. respond positive. For potassium hydroxide (KOH) test, bacterial isolates 4a2, JHL-12, 5C and RH-87 showed positive test results while other isolates were negative for KOH test. All the tested bacterial isolates were catalase positive. Many studies have reported catalase production by rhizobacteria (Ali Kamboh et al., 2009; Kumari et al., 2018). Catalase positive were reported to suppress early blight disease in tomato (Senthilraja et al., 2013) and induced resistance against tomato yellow leaf curl virus (Li et al., 2016). Out of 8 bacterial isolates, 1D and RH-87 were positive for levan production and three bacterial isolates JHL-12, 1C2 and 1D were fermenting the carbohydrates. Except RH-87, all the bacterial isolates exhibited positive response for oxidase test and RH-24 was negative for oxidative fermentative test. All the tested bacterial isolates showed nitrate reduction except 4a2 and RH-87. All the bacterial isolates showed gelatin hydrolysis activity except 4a2.

The 16S rRNA sequences have been widely used in the classification and identification of Bacteria and Archaea (Goodfellow et al., 2014). The sequence analysis of bacterial isolates depicted that tested antagonistic bacteria were belonged to three different genera including Flavobacterium spp. 4a2, *Bacillus* spp. JHL-8, 1C2, RH-24, 1D, *Pseudomonas* spp. JHL-12, 5C and RH-87. In our studies, 16S rRNA sequence based neighbor-Joining (N-J) tree indicated that two bacterial isolates ID (MH393211) and 1C2 (MH393210) showed 97 to 98% sequence identity with *Bacillus cereus* (accessions MK606105 and MK648339) while JHL-8 (MH393209) showed 99% sequence homology with *B. megaterium* (accession MG430236). The sequence of RH-24 (MH393208) had 99% sequence homology with *B. subtilis* (accession KY000519) and RH-87 (accession MT421780) showed identity 99% with *Pseudomonas libanensis* (DQ095905). Two bacterial isolates 5C (MH371201) and JHL-12 (MH371200) had 99% sequence homology with P. putida (accessions KY982927 and MF276642 respectively). Bacterial isolate 4a2 (accession MT421823) had 99% identity with the *Flavobacterium* spp. (accession HM745136). Similar 16S rRNA gene sequence based rhizobacterial characterizations have been reported in literature (Islam et al., 2016; Kuan et al., 2016; Kumari et al., 2018; Zouaoui et al., 2019).

Bacterial strains were also tested for plant growth promoting traits. In this study, all the antagonistic bacterial produced ammonia (NH_3_). Ammonia production is the most common character of PGPR, which indirectly enhances the plant growth (Yadav et al., 2010). It accumulates and supply nitrogen to their host plants and helps in plant growth promotion (Kumar et al., 2016), and it also contributes in antagonism (Howell, 1988). Various researchers have cited the ammonia production by rhizobacteria (Jayasinghearachchi and Seneviratne, 2004; Mazumdar et al., 2019; Triveni et al., 2012). Furthermore, all the tested bacterial isolates except 4a2 where able to hydrolyze starch. The production of HCN by PGPR is independent of their genus. Except 1D and 4a2, all the bacterial strains were positive for HCN production test. Previous researches have documented that the bacterial agents with HCN producing ability can be used as biocontrol agents (Ramette et al., 2003). Now, it is believed that HCN production indirectly increases phosphorus availability by chelation and sequestration of metals, and indirectly increases the nutrient availability to the rhizobacteria and host plants (Rijavec and Lapanje, 2016), thus they are used as biofertilizers or bio-control to enhance crop production (Agbodjato et al., 2015).

All the bacterial strains were also evaluated for P-solublization, indole-3-acetic acid (IAA) and siderophore production. Phosphate solubilization is an important plant growth promotion trait of rhizobacteria; in which rhizobacteria produce low molecular weight organic acids which solubilize phosphate thus, lowering the pH of the soil (Khan et al., 2014) thus, converts the phosphate into available forms that is taken up by the plant roots (Ahmad et al., 2018). In this study, all the bacterial isolates belonging to *Flavobacterium, Bacillus* and *Pseudomonas* spp. showed P-solublization ability and the test was not performed for 1D. Indole-3-acetic acid (IAA) is a vital phytohormone which is involved in in root development, elongation, proliferation and facilitates plants to obtain water and nutrients from soil (Yao et al., 2008). It increases the root surface area and loosens the plant cell walls, which facilitates in getting soil nutrients and supports better plant microbe interaction (Glick, 2012). In our study, all the bacterial isolates produced considerable amount of IAA (13.4–39.0 μgml^−1^) which were comparable to the previously published reports (Islam et al., 2016; Zahid et al., 2015). The increase in shoot and root length in bacterial inoculated plants may be attributed to their IAA production ability. Siderophore production is one of the most influential traits exhibited by the plant growth promoting rhizobacteria especially, when iron availability is limited (Whipps, 2001) and suppresses the phytopathogens by depriving them of available iron (O’sullivan and O’Gara, 1992). All the tested PGPR tested in this study showed promising siderophore production (17.1– 27.7%) which proofs rhizobacterial ability to suppress the growth of target pathogen *P. myriotylum*. Many previous reports have supported siderophore production potential of PGPR and their role in disease suppression (Ali et al., 2020; Compant et al., 2005; Sayyed et al., 2019; Swadling and Jeffries, 1996).

Previous studies have proven that heavy metal ions co-regulate genes which confer antibiotic resistance and decrease antibiotic susceptibility (Baker-Austin et al., 2006; Rani et al., 2010). In this study, bacterial isolates were screened for their resistance or susceptibility response against Streptomyces, ampicillin, rifampicin, penicillin G and vancomycin at five dose levels, and all the tested bacterial strains displayed varied level of resistance and antibiotic susceptibility response. The study has shown that the tested bacterial isolates have varied levels of tendencies to overcome the antibiotics stress and it might be associated with tolerance against heavy metals present in soil. Bacterial isolates showing multiple antibiotic resistances have greater chances to establish as inoculum in the soil and any new niche. Metal tolerance and antibiotic resistance have previously been reported (Thacker et al., 2007; Wani and Irene, 2014).

Treatment of chilli seed with selected bacterial strains significantly improved the seed germination, plumule and radical length and vigor index as compared to un-inoculated control. No phytotoxicity and stress on seedling germination had been observed in any treatment. All the antagonists used in this study were non-pathogenic to chilli seeds. As these bacterial isolates showed P-solubilization and IAA activity thus, can be used for plant growth promotion (Naureen et al., 2009) and the growth enhancement may be due to the production of IAA. A research study has shown high amylase activity during seed germination in rice and legume inoculated with PGPR (Duarah et al., 2011).

The level of defense-related enzymes contribute significantly in the mechanism of host plant resistance (Shivakumar et al., 2000). The bacterial inoculated chilli plants also displayed a significant increase in the defense related enzymes activities. Chilli seedlings treated with bacterial isolates exhibited significant increase in Peroxidase (PO), Polyphenol oxidase (PPO) and Phenylalanine ammonialyas (PAL) activities. PO activity was maximum in the seedlings treated with 4a2 followed by JHL-8 and RH-24 while PPO and PAL activity was recorded significantly high 5 days after inoculation (DAI) in chilli roots bacterized with RH-24 inoculum over un-inoculated (UC) and negative control (NC). Our results are supported by the findings (Benhamou and Paulitz, 2000) where cucumber roots inoculation with *Pseudomonas corrugate* and *P. aureofaciens* suppressed the root and crown rot caused by *P. aphanidermatum* and PAL accumulation lasted for 16 days while Peroxidase (PO) and Polyphenol oxidase (PPO) activities were enhanced in roots 2 to 5 days after bacterial treatment. It was previously reported that the induction of plant defense related enzymes is related with the plant defense system and induced resistance by PGPR inoculation and colonization (Liang et al., 2011). Accumulation of defense related enzymes after PGPR application has been reported in cucumber (Liang et al., 2011), chilli (Jayapala et al., 2019) and tomato (Ramamoorthy et al., 2002).

Many studies have proved that PGPR have great potential to suppress the plant pathogens and increasing plant growth under greenhouse conditions (Kabdwal et al., 2019). These PGPR induce systemic resistance against a broad spectrum of pathogens due to their root colonization ability (Salem and Abd El-Shafea, 2018). Both *Pseudomonas* and *Bacillus* spp. are known for their role in disease suppression against various plant pathogens (Velusamy et al., 2013). In present studies, tested bacterial strains enhanced the chilli germination percentage and reduced the seedling mortality percentage by suppressing the *P. myriotylum*, and a significant increase in plant growth characters was observed in bacterial inoculated treatments over un-treated (UTC) and negative control (NC). Similar results have been reported in other research studies (Almaghrabi et al., 2013; Egamberdieva, 2011; Islam et al., 2016; Torres et al., 2020).

Many researches have disclosed the low performance of the bacterial based products under open field conditions due to various climatic and soil factors (Ownley et al., 2003) which affect the bacterial colonization ability, biological activates, and disease suppressing potential (Landa et al., 2001). Biocontrol is a complex phenomenon involving several mechanisms in disease suppression and understanding these mechanisms would be beneficial for the effective utilization of bacterial biocontrol agents in open fields. Many biocontrol products against damping-off disease are available worldwide (Lamichhane et al., 2017) but not a single locally prepared product is available and registered in Pakistan. Bio-products imported from other countries failed to perfume due to different soil nature and climatic conditions. Keeping in view these facts, this study was performed to screen out the native bacterial antagonists with high disease suppressive and plant growth promotion ability. To our knowledge antagonistic potential of native PGPR was first time reported from Pakistan. However, series of experiments are required to test the efficacy of these bacterial isolates at different dose levels and formulations with different soil types and climatic conditions. Finally, field trials will help to develop the bacterial based bioproduct, its registration and commercialization.

## CONCLUSION

Plant disease suppression and growth promotion are considerable aspects to get good quality produce. Chilli pepper is cultivated as an important vegetable crop across the world, and its production is greatly reduced by damping-off disease caused by *Pythium myriotylum.* Having this in mind, we screened rhizobacterial isolates in vitro which showed greater potential to suppress *P. myriotylum* inoculum, and significantly improved the seedling germination and vigor index without posing any phytotoxicity and pathogenic impact on young seedlings. These bacterial isolates showed P-solubilization, indole-3-acetic acid and siderophores production ability and produced varied levels of susceptibility and tolerance response against antibiotics. The application of bacterial isolates increased the accumulation and activities of defense related enzymes (PO, PPO and PAL) in chilli roots under pathogen pressure. These PGPR having multiple disease suppressive and PGP traits significantly reduced the seedling mortality and improved the seedling germination and other PGP traits in pot trials. However, detailed studies about dose calibration and field performances are required to ensure safe application of these bacteria.

## Acknowledgments

Financial support received from Higher Education Commission (HEC) and Punjab Agriculture Research Board (PARB), Pakistan for carrying out this research work is gratefully acknowledged. We also acknowledge Professor Youfu Frank Zhao, Department of Crop Sciences, University of Illinois at Urbana-Champaign for providing bench space in his laboratory and helping in various research activities.

## AUTHOR’S CONTRIBUTIONS

**Sajjad Hyder**

Designed the research, conducted the experiments and wrote the manuscript

**Amjad Shahzad Gondal**

Helped in data analysis and write up and proofreading

**Zarrin Fatima Rizvi**

Helped in data analysis and proofreading

**Muhammad Irtaza Sajjad Haider**

Helped in research work and data collection

**Muhammad Insam-ul-Haq**

Supervised the research work

